# Structural Insights into γH2Ax containing Nucleosomes

**DOI:** 10.1101/2023.04.30.538894

**Authors:** Rashmi Panigrahi, Ross Edwards, Md Touhidul (Apu) Islam, Jun Lu, Ayodeji Kulepa, Tae Hwan Kim, J. N. Mark Glover

## Abstract

MDC1 is a key mediator of DNA-damage signaling. When DNA double-strand breaks (DSB) occur, the histone variant H2AX on the nucleosome is phosphorylated on its C-terminus at residue Ser139 to form the γH2AX nucleosome. This phosphorylated form is specifically recognized by the tandem BRCT repeats of MDC1. The MDC1-bound nucleosome serves as a docking platform to promote the localization of other DNA repair factors. To further characterize the nucleosome-BRCT interaction, we developed a time efficient two-step modified native chemical ligation protocol to prepare phosphorylated nucleosomes. Our binding studies show that BRCT interacts with the nucleosome with a higher affinity than the phosphorylated peptide. Using cryogenic electron microscopy (cryo-EM), we obtained structures of the γH2AX nucleosome revealing the structural basis for nucleosome-nucleosome stacking promoted by interactions of the H4 N-terminal of one nucleosome with its stacked partner. In contrast, we show that binding of the MDC1 BRCT domain disrupts this stacking, suggesting that histone/DNA dynamics are integral to DNA damage signaling.

## Introduction

The nucleosome is the fundamental repeating element of chromatin and is crucial for DNA packing. There is a five-fold contraction in the length of DNA in the nucleosome and a 10,000-fold contraction in DNA length in the condensed states of mammalian chromosomes[1]. Nucleosomes regulate accessibility of the DNA to other DNA binding proteins, which is at the core of gene regulatory processes. Decades of accumulating evidence have reiterated the importance of histone post translational modifications in the signaling of various regulatory processes. Here we focus on the signaling that elicits DNA repair. An approximately 1.6 left-handed superhelical turns of 145-base pair (bp) double stranded DNA (dsDNA) wraps around an octameric core of four histone proteins, H2A, H2B, H3 and H4 to form a nucleosome. This octamer contains two copies of histone H2A-H2B dimers and one copy of an H3-H4 tetramer. Decades of accumulating evidence have demonstrated that chromatin structure plays a crucial role in gene expression by regulating DNA accessibility through histone posttranslational modifications (PTM). The PTMs were annotated since 1960s[2, 3] and the development of genome-wide mapping approaches and high-resolution structure determination methods bolstered our understanding of their direct and indirect effects on the genome function thus underlining their significance. These PTMs include phosphorylation, acetylation, ubiquitination, methylation, to name a few, and are present both in the histone core and their N- and C-terminal tails[3-5]. These PTMs also known as histone codes are indispensable epigenetic marks[6]. At a fundamental level, PTMs modify the nucleosome to recruit or discharge specific regulatory partners by altering their affinities for the nucleosome [4]. The PTMs can either directly drive conformational changes in the nucleosome by modifying histone-DNA interactions or the PTMs can be recognized by the mediator proteins or readers, which in turn recruit a battery of proteins in a concerted manner to the nucleosome[3, 5]. This works focuses on the phosphorylation of H2AX, which acts as a signal for DNA damage response.

DNA damage is an unavoidable event in living cell as the genome is constantly exposed to environmental and endogenous genotoxic agents, creating thousands of lesions per day. Of these lesions, DNA double-strand breaks (DSBs) are the most problematic for genome stability. To handle these lesions, cells have evolved intricate DNA repair pathways to neutralize these genomic insults. DSBs can be repaired either via the homologous recombination (HR) or the nonhomologous end-joining (NHEJ) pathways. Immediately after DSB induction, the H2AX variant that comprises 10–15% of total cellular H2A, is phosphorylated at Ser^139^ generating γH2AX foci. [7-9]. This phosphorylation acts as an epigenetic mark that flags megabase regions around individual break and is an essential and efficient coordinator of DSB recognition and DNA repair[10]. The H2AX gene, a member of the H2A family, is located at 11q23.2-q23.3 and encodes a 142 amino acid protein. The histone H2AX contains 143 amino acids while the H2A contains 130 residues. The residues of C-terminal tail after residue 122 are highly variable between the two histone variants (Extended Data Fig. 1A). The C-terminus has the consensus Ser-Gln-Glu (SQE) motif which is a common recognition site for phosphorylation by the phosphatidylinositol-3-OH-kinase-like family of protein kinases (PIKKs) such as ATM (ataxia telangiectasia mutated)[11], ATR (ATM and Rad3-related)[12, 13], or DNA-dependent protein kinase (DNA-PK)[9, 14]. The γH2AX foci specifically attracts repair factors in a coordinated fashion. The BRCA1 carboxy-terminal (BRCT) repeats of the mediator of DNA damage checkpoint protein 1 (MDC1) is the predominant recognition module of γH2AX[15-17]. MDC1 is a 2089 amino acid containing protein with two phosphorylation specific interaction modules: the N-terminal forkhead-associated (FHA) domain and the C-terminal BRCT domain. These two domains are linked by a disordered region which contain multiple phosphorylation sites[10]. In this study, we sought set out to decipher the interaction between the MDC1 BRCT domain and γH2AX containing nucleosome.

First, we developed a time efficient one-step purification method to produced human histone octamer lacking the C-terminal region “ATQASQEY” of H2AX. Next, we used native chemical ligation- to ligate the phosphorylated Ser^139^ (pSer^139^) containing peptide to the octamer. The native chemically ligated octamer was assembled with 145 bp DNA to obtain γH2AX containing nucleosomes. Electrophoretic mobility shift assay (EMSA) and microscale thermophoresis (MST) confirmed the interaction of human MDC1 BRCT (hMDC1) with the phosphorylated nucleosomes. Cryo-EM was used to decipher the structure of the γH2AX containing nucleosomes. A novel stacking interaction was observed in the nucleosome alone which was broken in the presence of hMDC1 BRCT. With this observation we propose that BRCT domain acts as a reader of phosphorylation of the γH2AX nucleosomes. BRCT binding to the phosphorylated S139 on γH2AX impedes nucleosome stacking interaction.

## Results

### Two step purification of the γH2AX containing histone octamer

Different methods of histone purification have been established to date including the refolding method[18], histone chaperone mediated co-expression[19] and polycistronic expression[20]. In order to study histone PTMs two methods have been largely used, an E. coli cell-free protein synthesis system[21, 22] and native chemical ligation of single histones purified using the denaturation method. We have developed a modified native chemical ligation (NCL) method, which does not require denaturation of histones thereby avoiding protein loss due to precipitation during refolding. Our co-expression system takes advantage of the duet vector system, which allows the four histones to be co-expressed in E. coli. After purification of the octamer using Ni-NTA affinity chromatography, the γH2AX peptide is attached to the C-terminus of H2AX using the NCL method (Fig. 1A). The presence of pSer^139^ was confirmed by western blot using γH2AX specific antibody (Fig. 1B). The phosphorylated octamer was further purified using size exclusion chromatography (Extended Data Fig. 2A). For the control experiments, the unmodified H2AX containing octamer was purified using co-expression method (Extended Data Fig. 1B). After the successful nucleosome assembly, both unmodified nucleosome and γH2AX containing nucleosome were tested for their binding with hMDC1 BRCT. The BRCT showed interaction specifically with the phosphorylated version (Fig. 2A). These findings agree with the previous findings from the high-resolution crystal structures where a peptide mimicking the γH2AX C-terminus were used to observe the interactions[16, 17]. Fluorescence polarization (FP) studies indicated that the phosphorylated C-terminal tail peptide interacts with the BRCT with 1.6 μM affinity (Extended Data Fig. 2B). MST experiments did not indicate any interaction between the MDC1 BRCT and the unmodified nucleosome, however a tight interaction with a binding affinity of 14.2 nM is observed with the γH2AX nucleosome indicating that phosphorylation of H2AX is critical for BRCT binding. Our findings indicate that the hMDC1 BRCT interact with the nucleosome primarily through the pSer^139^-containing C-terminal tail.

**Figure 1.**
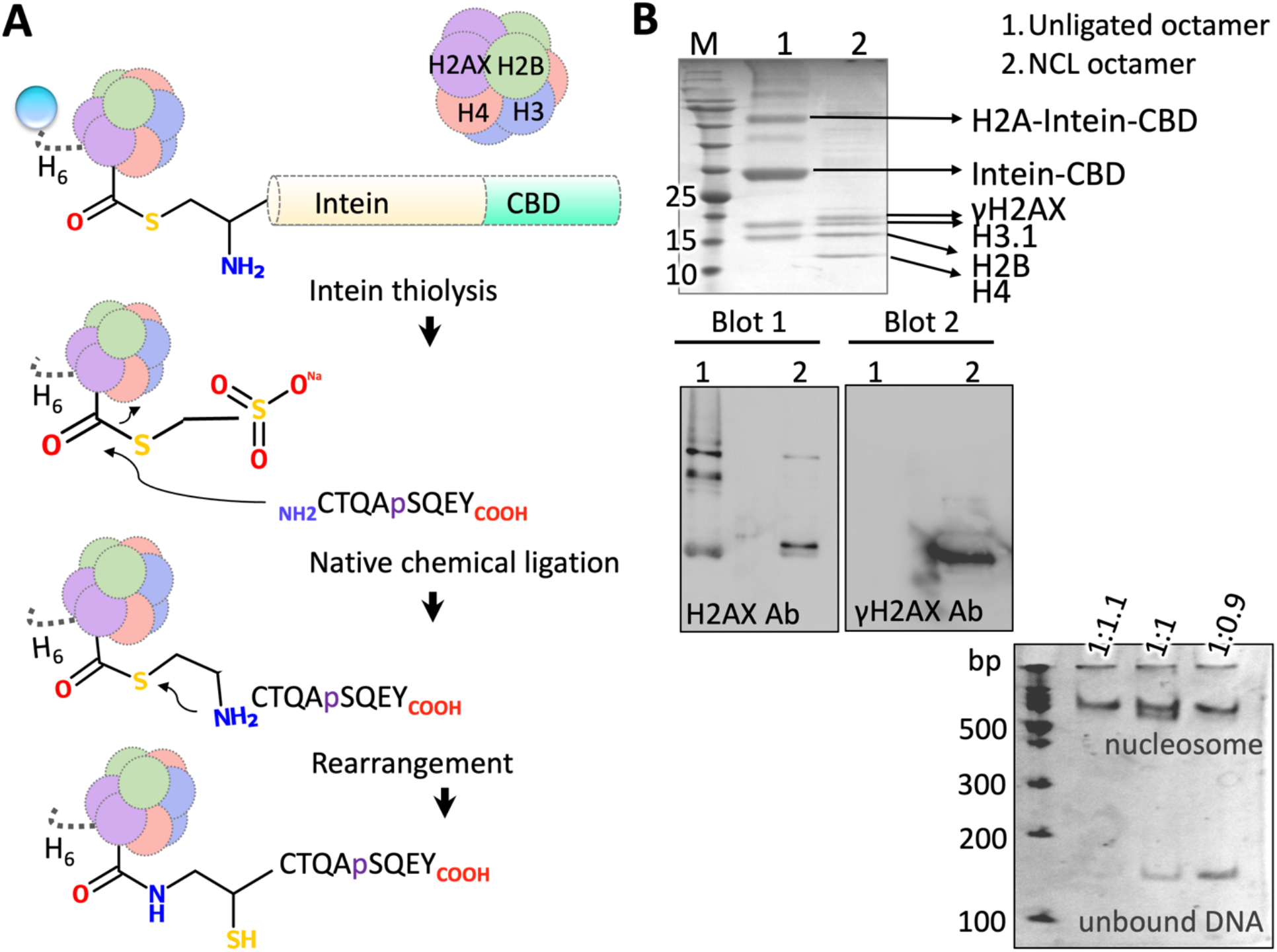
Native chemical ligation for γH2AX octamer preparation. **A**. Step by step protocol. The his-tagged octamer is subjected to intein thiolysis by addition of MESNa. Phosphpeptide is added which initiates the NCL process. **B**. SDS PAGE showing different histones where lane 1 represents unligated octamer and lane 2 represents ligated octamer. Molecular weight marker (M) is shown in kDa. The center panel shows western blot with anti-H2AX antibody (left) and anti-γH2AX antibody (right). The western blot (right) confirms NCL is successful and that γH2AX is present only in the lane containing ligated octamer. The γH2AX containing nucleosome was used for nucleosome assembly (bottom panel). At the ratio of octamer to DNA (1.1:1) no unbound DNA is observed.

**Figure 2.**
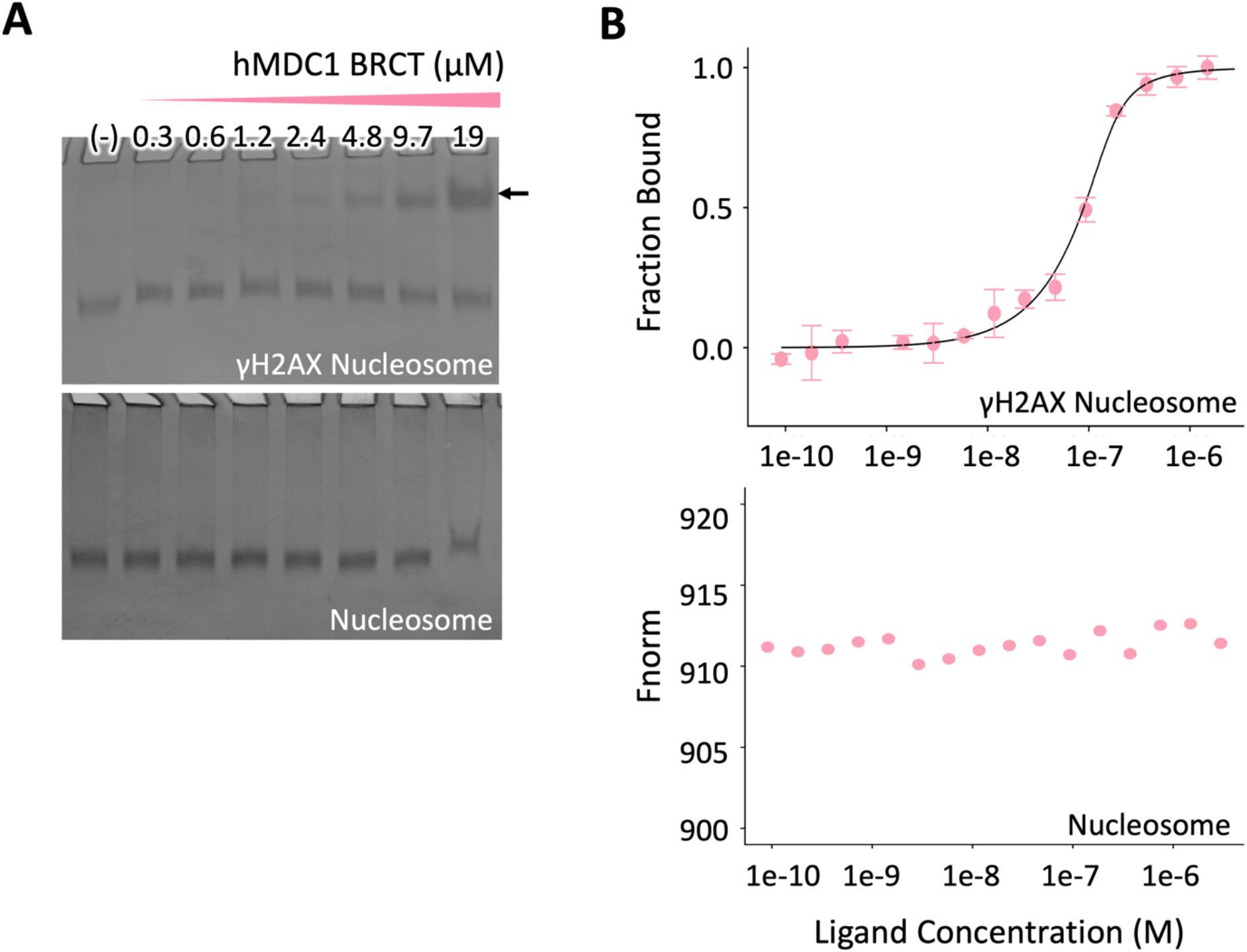
Binding studies of hMDC1 BRCT with γH2AX containing nucleosome. **A**. EMSA studies **B**. MST studies. The control runs were performed to test the interaction of hMDC1 BRCT with nucleosome. The EMSA gels are stained for DNA using SYBR Safe.

### Structure of the canonical γH2AX containing nucleosome

The cryo-EM data was collected for both unmodified and γH2AX containing nucleosomes. The particles were arranged as arrays of stack with a preferred orientation (side views) under standard vitrification conditions (Extended Data Fig. 3). In order to get a closer look at the stack interface, we collected cryo-EM data at different tilt angles to capture structural details at different sample orientations. The structure of unmodified nucleosome is identical to its phosphorylated counterpart (Extended Data Fig. 1C). Here we report the structural finding pertaining to the γH2AX containing nucleosome (Extended Data Table S1). A data set of 2,512,360 particles was collected, and several rounds of 2D and 3D classifications revealed views of mononucleosomes in a set containing 127,051 particles with different orientations: as a disk, tilted views, and side views (Supplemental Table S1, Fig. 3A, Extended Data Fig. 4). Structural details such as the DNA dyad, major and minor DNA grooves and histone α-helices can be clearly seen, and a final map was calculated to a nominal average resolution of 3.4 Å (Fig. 3A, 4 and Extended Data Fig. 5A). The cryo-EM structure of the Xenopus nucleosome (PDB: 7OHC) determined at 2.5 Å was used as the reference model for model building and refinement[23]. Two copies each of the four core histones (H3, H4, H2A and H2B) is wrapped by the left-handed super helical 601 – 145 bp dsDNA (Fig. 3A, 5A). The local resolution for the histone core is higher than the DNA (Fig. 3A and 4). A slight increase in the nucleosome size was noticed in our canonical structure when compared to the nucleosome from the Xenopus (Fig. 5). The location where the major grooves face the histone octamer or the ‘superhelix location’ (SHL), was used to mark the differences in DNA positioning between our canonical γH2AX nucleosome and the reference model. The γH2AX nucleosome expands in the direction perpendicular to the symmetry axis or the dyad axis compared to the reference model (Fig. 5). Although there is no change in DNA at SHL 0 and 4, the difference between the two structures become clear at SHL positions 2, 3, 5 and 6. The height of the nucleosome is 55 Å and there is no change in the distance between the two DNA gyres from the side view (Fig. 5A), indicating a more lateral dilation, in case of co-expressed γH2AX nucleosome when compared to the reference model. Similarly, there is no widening of the DNA major/minor grooves, which are otherwise observed upon binding of certain transcription pioneer factors like SOX11[24]. During the dimerization of H2AX – H2B and H3.1 – H4 pairs, the loop L1 of one histone aligns with the loop L2 of the other forming the handshake motif. This handshake motif interacts with DNA via two L1L2 sites and one α1α1 site. Rearrangement in the histone secondary structures is observed along the direction of motion of DNA when compared to the reference model (Fig. 5B and 6). In the histone γH2AX, the motion in DNA at SHL 5.5 drags the L2 loop, which in turn causes an upward motion in the α2 (Fig. 6A). Similar concerted motion with DNA – α1 – L1 – α2 was observed in H2B (Fig. 6C). In the case of the histone H3.1, the motion is primarily observed in the region which is in contact with DNA including αN, α1, L1, α2 (Fig. 6B). In the case of histone H4, the motion is primarily observed in α1, L1 and α2 (Fig. 6D). The flexible tails of the core histones interact with DNA via the minor groove. For the γH2AX C-terminal tail, electron density could be traced until Lys^119^, however there was no density visible for the remaining 23 residues. Most of the important histone-DNA contacts are previous described are observed. For instance, residues Arg^43^, Gly^45^, Val^47^, Arg^64^, Arg^70^, Arg^73^, Arg^84^, Ser^87^, Val^118^, Thr119 from histone H3.1; residues Arg^36^, Arg^37^, Ile^47^, Gly^49^, Lys^80^, Thr^81^ from histone H4; residues Arg^12^, Lys^14^, Arg^18^, Arg^30^, Arg^33^, Val^44^, Ala^46^, Thr^77^ from histone γH2AX and residues Arg^31^, Lys^32^, Ser^54^, Arg^84^, Ser^85^, Thr^86^ from histone H2B, hold the 601 DNA in place to form a canonical nucleosome. The resolution of 601 DNA is lower than the histone core reflecting DNA flexibility (Fig. 3A bottom panel).

**Figure 3.**
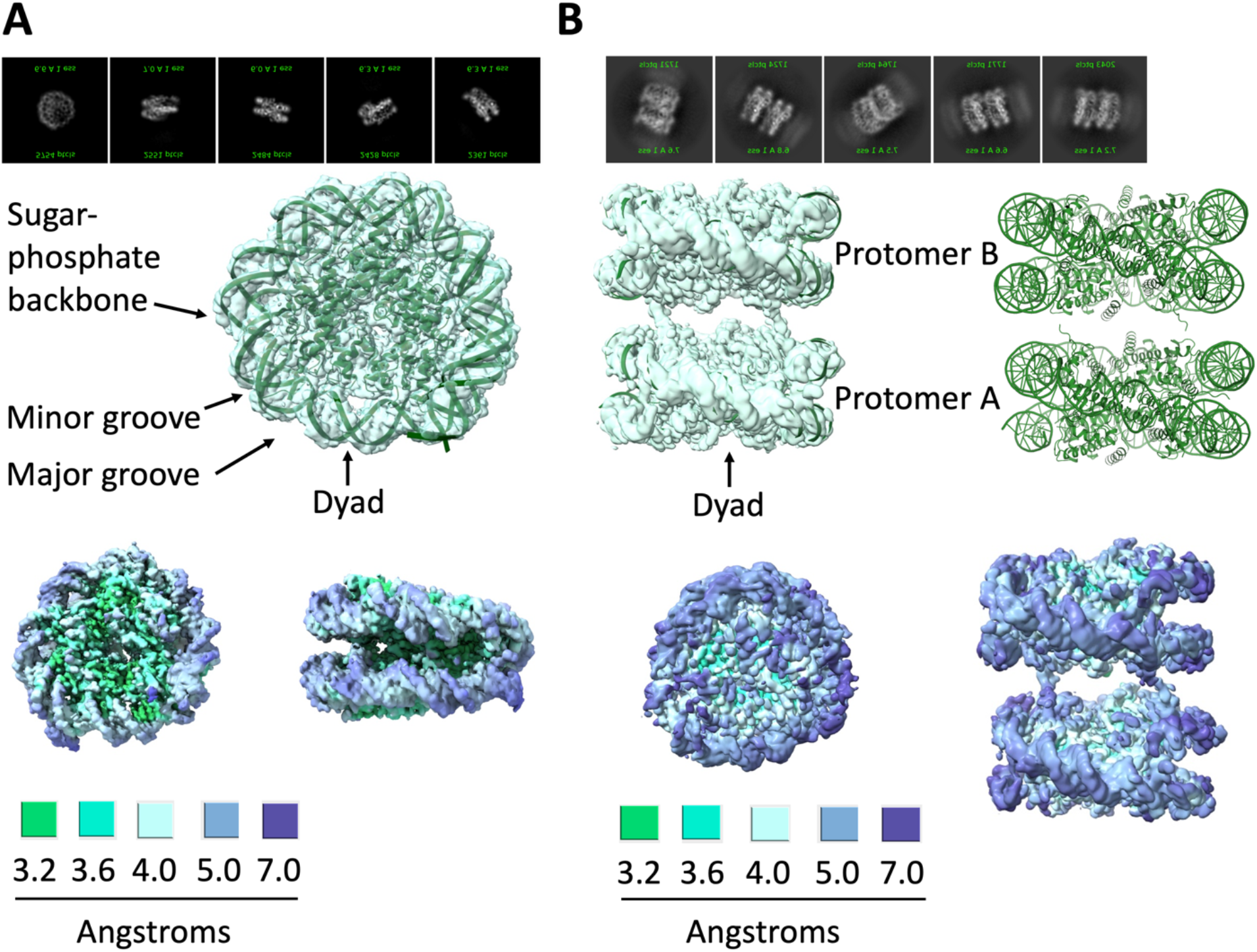
Cryo-EM structures of canonical γH2AX nucleosome and γH2AX nucleosome stack. **A**. Canonical nucleosome map with model in surface and cartoon representation respectively, showing defined sugar-phosphate backbone, major and minor grooves and dyad. The top panel shows the different orientations captured during 2D classification. **B**. Stack nucleosome. The two protomers and the location of dyad is labeled. The top panel shows the snapshot of the different orientations captured during 2D classification. Bottom panels show the local resolution maps and the areas with certain values for resolution are color coded.

**Figure 4.**
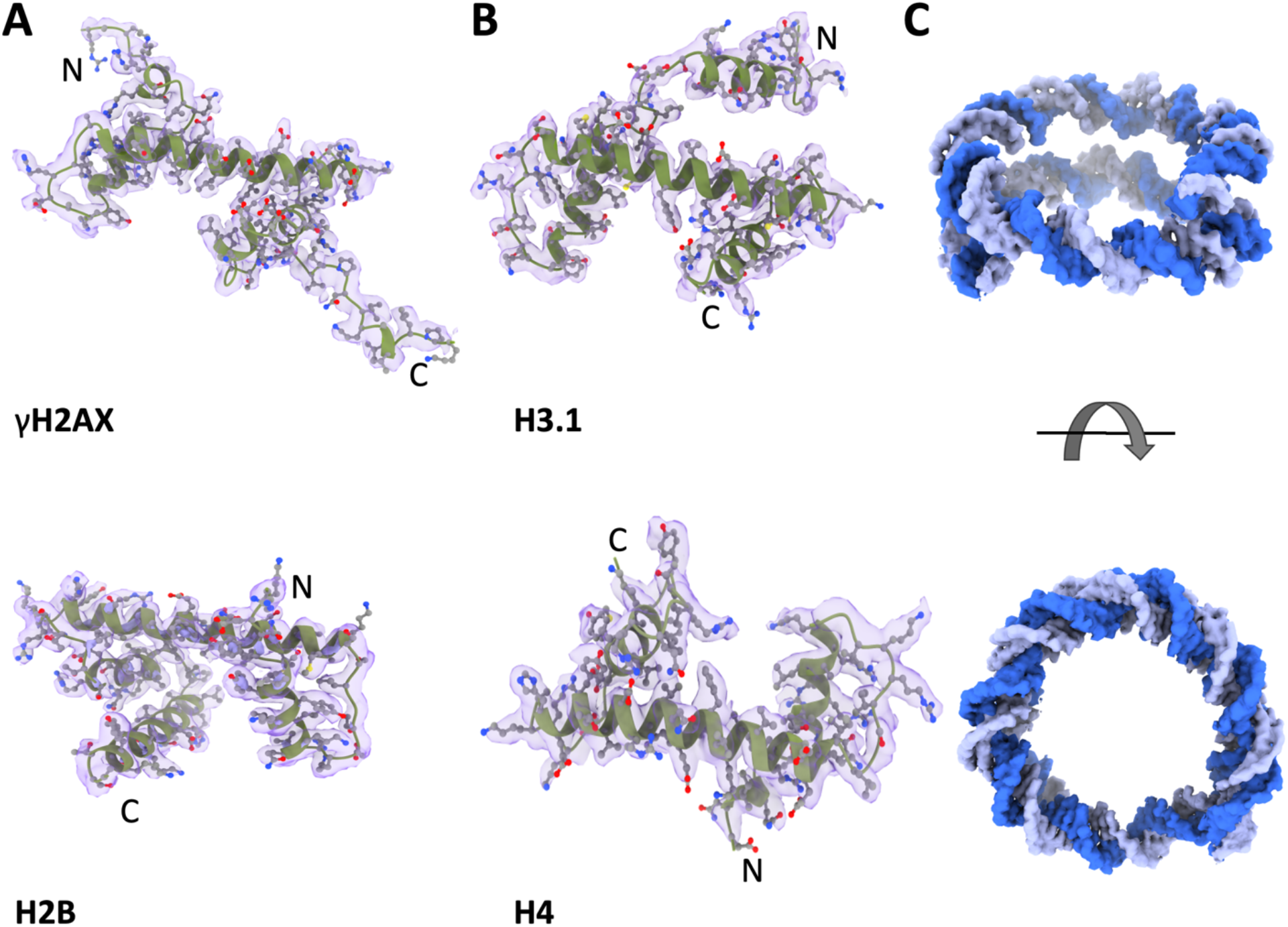
Quality of the cryo-EM map and the model for the canonical γH2AX nucleosome structure. Density map and the fitted models corresponding to the four histones γH2AX (**A** top panel), H2B (**A** bottom panel), H3.1 (**B** top panel) and H4 (**B** bottom panel) are shown. Surface representation of 601 dsDNA is shown in panel **C**.

**Figure 5.**
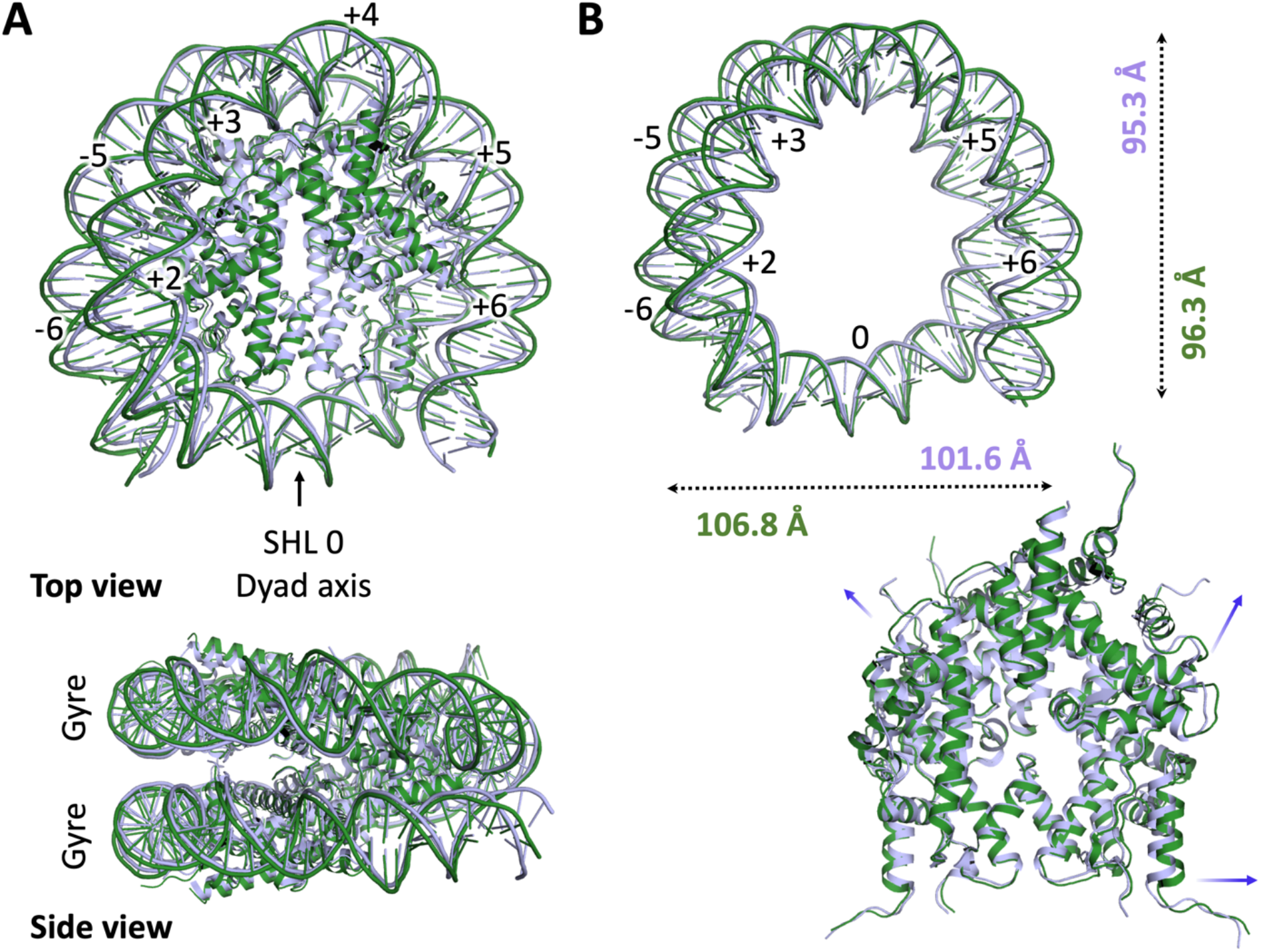
Canonical γH2AX nucleosome structure in comparison to the reference model (PDB: 7OHC). **A**. Top view of canonical γH2AX nucleosome (forest green) overlayed on xenopus nucleosome (light blue) obtained using cryo-EM Location of dyad axis and numbers indicating SHL positions are shown. The bottom panel shows the side view of the overlay, indicating that there is no change in the position of the two DNA gyres. **B**. Change in the DNA position at certain SHLs from the top view. The nucleosome expansion in the direction perpendicular to the symmetry axis (distance between C3 in nucleotides -19 and -19) and along the symmetry axis (distance between C3 in nucleotides 2 and 38) is shown. The values are color coded to match with the respective structures. The bottom panel shows motion in the histones’ secondary structures which is correlated with that of the DNA double helix.

**Figure 6.**
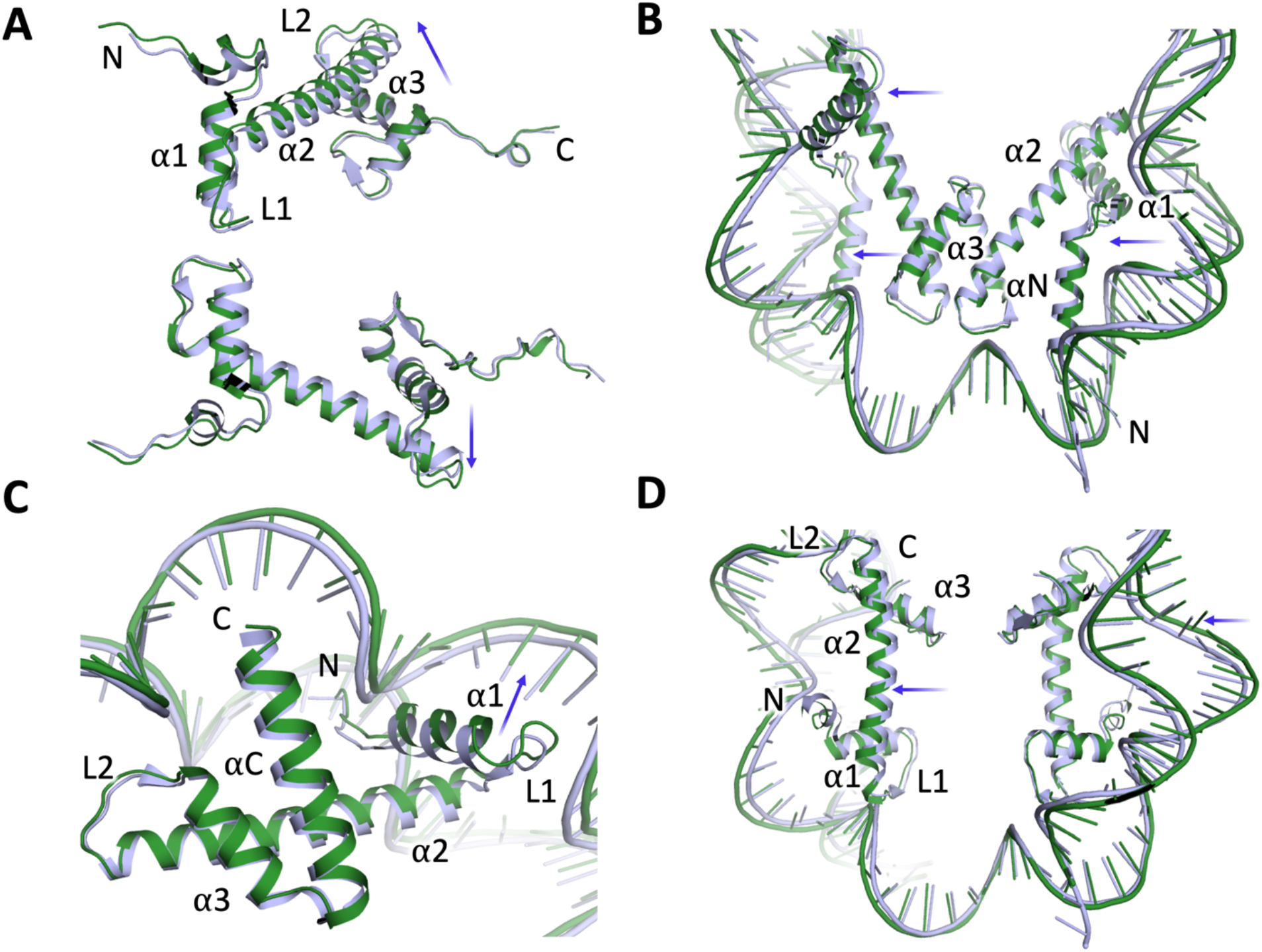
Comparative motion between the canonical γH2AX nucleosome (forest green) and the reference model in light blue (PDB: 7OHC). All the secondary structural elements are shown for **A**. H2AX; **B**. H3.1; **C**. H2B; and **D**. H4. The motion of the histone secondary structural elements (α for helix and L for loop) with respective DNA are shown with blue arrows. One histone protomer is labeled and single DNA strands for both structures are shown for clarity.

### Structure of parallel stacked γH2AX-containing nucleosomes

Our cryo-EM grids showed arrays of nucleosomes stacks, potentially representing a 10 nm chromatin fiber like arrangement (Extended Data Fig. 3). A class containing 118,190 particles were used to build the stacked nucleosome structure determined at 4 Å resolution (Supplemental Table S1, Extended Data Fig. 4) and comprised of two nucleosomes stacked with their DNA entry/exit site facing the same direction (Fig. 3B). The two nucleosomes are arranged with their dyad axis parallel to each other and the distance between their SHL 0 positions is ∼61 Å. Our canonical γH2AX nucleosome model fits well into the densities of each protomer in the stack. Clear main chain densities for H4 N-terminal tail from one nucleosome interact with stacked partner nucleosome (Fig. 7). Five residues of the H4 N-terminal tail, Arg^20^, Lys^21^, Val^22^, Leu^23^ and Arg^24^, could be built into this density using ISOLDE (Fig. 7). The sidechain of Arg^24^ lacks clear density and is likely flexible. Arg^20^ of one nucleosome is positioned near the adjacent nucleosome of the stack via hydrogen bonding interactions with Asp^49^ of H2B. The carboxylate group of Asp^49^ and the guanidinium of Arg^20^ form salt bridge that holds Arg^20^ in place and aids in the nucleosomes stacking. The side chain of Lys^21^ forms hydrogen bond with the phosphate backbone of the DNA of its own protomer. Similar to the canonical γH2AX nucleosome, the nucleosome stacks showed low resolution for DNA (Fig. 3B bottom panel). Histone dynamics mediated nucleosome unwrapping has been studied using cryo-EM, NMR[25] and molecular dynamics[26, 27]. The missing density for the DNA backbone may indicate dynamics (Extended Data Fig. 6).

**Figure 7.**
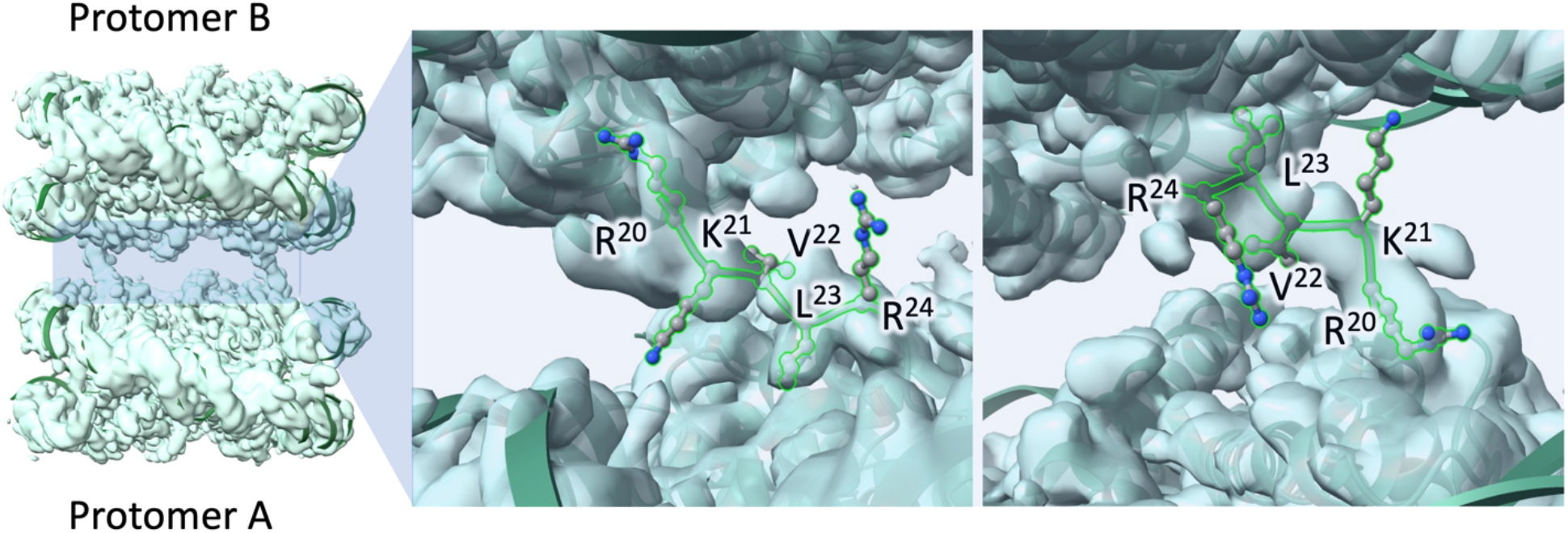
H4 tail interaction between protomer A and B in the γH2AX nucleosome stack structure. The residues of H4 tail involved in the inter-protomer interaction was build using ISOLDE.

We prepared cryo-EM samples of γH2AX nucleosome in complex with BRCT. Although no density for the BRCT was observed, the particle orientation captured on the grids showed a striking difference compared to the nucleosome only grids. The nucleosome particles were no longer arranged in a preferred orientation or stack like arrangement (Extended Data Fig. 3B) in presence of BRCT. The hMDC1 BRCT is 22 kDa and interacts with pSer^139^ and Tyr^142^ as evidenced by crystallographic studies[16, 17]. It is possible that this 23-residue stretch bearing the BRCT protruding out of the nucleosome disc from Lys^119^ at the C-terminal end of γH2AX remains disordered, and hence no density for BRCT was observed in the cryo-EM grids containing γH2AX nucleosome – BRCT complex.

## Conclusion

The packing of DNA into chromatin is essential for regulation of cellular process like transcription, epigenetic regulation, and DNA repair. At the heart of these processes, nucleosome, the basic unit of chromatin, play a dynamic role as revealed by high-resolution structural studies[28, 29]. Although histone-histone and histone-DNA interactions render nucleosome its stability, it is not a static disc. In the cryo-EM structure of γH2AX containing nucleosome, we observe a slightly wider nucleosome along the axis perpendicular to the dyad symmetry axis. This widening of the nucleosome accompanied with proportional motion of histone secondary structural elements, while the no change is observed at SHL 0 when compared to the reference model. Similar observation is noted in our unique parallelly stacked γH2AX nucleosome structure. Interphase chromosomes display an extensively condensed nucleosome array. Cryo-EM structure of 30 nm chromatin fiber designed using 12 repeats of 601 DNA in the presence of linker histone H1[30] show zigzag configuration of nucleosome arrangement. Similar findings were observed in the presence of linker histone H5 with low NaCl concentration supplemented with Mg^2+^ (1.6 mM – 5 mM)[31]. In the 30 nm fiber structure, the presence of linker histone imposes additional impact on the rotation and the specific distance between the two adjacent protomers of the stacks. Thus, a strong interaction between the H2B-helix α1/αC and the adjacent H2A-helix α2 is observed at the interface between the adjacent histones. In our cryo-EM structure of the stacked γH2AX nucleosomes, we observe clear main chain densities for the N-termini of the H4 tails. The importance of H4 tail in stabilizing stacking interactions had been predicted using both conventional and steered MD simulations[32, 33]. However, the fact that we observe stacking interactions in the absence of linker DNA/histones, suggests that our co-expression method of purifying intact recombinant octamer could assist in formation of such stable nucleosome array under cryogenic conditions, compared to the widely used refolding method.

Histone tails are sites of PTMs and these modifications change the properties of nucleosomes such as charge and rigidification which leads to DNA/histone dynamics thus changing the chromatin organization. These changes facilitate gene regulation and repair. We observed flexibility in DNA characterized by lower resolution compared to the core histones. The flexibility is more towards the DNA entry/exit site in the nucleosome stack compared to the canonical. This suggests that stacked nucleosomes can transition to mononucleosome structures on phosphorylation to facilitate the interaction with repair factors. Further our findings underscore that pSer^139^ signals the recruitment of hMDC1 BRCT and is solely responsible for the interaction of BRCT with nucleosome. The nucleosome lacking this signal does not interact with the BRCT.

## Methods

Cloning of histones and overexpression. Human histone (H2AX, H2B, H3.1 and H4) genes were amplified by PCR from the plasmids originally obtained from the RIKEN laboratory. A hexahistidine tag followed by a Tobacco etch virus (TEV) protease recognition site was added to the N-termini of both H2AX^(1-134)^ and H3.1^(1-136)^. The DNA sequence pertaining to Mxe intein-chitin-binding domain (CBD) was PCR amplified from pTXB1 (New England Biolabs) and ligated in frame at the C-terminus of H2AX^(1-134)^. The PCR amplified products pertaining to the histone pairs, H2AX^(1-134-intein-CBD)^ – H2B^(1-126)^ and H3.1^(1-136)^ – H4^(1-103)^ were clone into duet vectors. The clones are available at addgene (Plasmid# 197272, Plasmid# 197273). The resulting plasmids encoding histone pairs was transformed into BL21(DE3)pLysS cells and plated on to an LB agar plate containing ampicillin (50 mg/mL) and chloramphenicol (25 mg/mL). The plate was incubated overnight (∼18 h) at 37 °C. For a 2 L preparation, a starter culture was prepared where one colony was inoculated into 100 mL media containing the above antibiotics. After an overnight growth at 37 °C, the starter culture was added to the autoclave sterilized 2 L media and was shaken at 160 rpm. When the OD600 reaches ∼0.6 – 0.8, histone co-expression was induced by adding 1 mM isopropyl β-D-1-thiogalactopyranoside (IPTG). The culture was grown for 6 hours at 30 °C. Cells were harvested by centrifugation at 4,500 rpm for 20 min at 4 °C. Cell pellets were processed immediately or stored at -80 °C for future purification.

### Two-step purification of histones

A nickel affinity chromatography step followed by a size exclusion chromatography was used to produced native chemically ligated γH2AX containing octamer. Cell pellets were resuspended in 100 ml of lysis buffer (20 mM Tris-HCl pH 7.0, 2.0 M sodium chloride (NaCl), 1 mM phenylmethanesulfonylfluoride. Resuspended cells were lysed by EmulsiFlex-C3 high pressure homogenizer (Avestin) and clarified by centrifugation at 12,000 rpm at 4 °C for 1 h. The clarified supernatant was incubated with Ni-NTA agarose beads (Qiagen) for 1 h and was applied on a gravity flow column to remove the bacterial protein contaminants. The beads were washed with lysis buffer. γH2AX was synthesized using a ligation-desulfurization strategy[34] (Fig 1A), where the C^α^-carboxy thioester of H2AX^(1–134)^ was first prepared by thiolysis of H2AX^(1-134-intein-CBD)^ fusion protein with MESNa (sodium 2-mercaptoethanesulfonate) via intein-mediated protein splicing. 25 mM MESNa was added to the beads in the gravity column and incubated for overnight at room temperature. The flow through containing intein-CBD was discarded. The octamer containing MES (2-mercaptoethanesulfonate) thioester of H2AX^(1–134)^ was eluted using imidazole gradient of 250 mM, 500 mM and 1 M. The MES thioester of H2AX^(1–134)^ containing octamer was then ligated with the synthetic C-terminal octapeptide containing the phosphorylated-Ser residue, ^NH2^-CTQApSQEY-^COOH^, to form the full-length γH2AX containing an Ala-Cys mutation at position 135, H2AX (A135C, pS139). The ligation was initiated by adding 300 mM TCEP (final concentration) to the octamer – peptide mixture. Desulfurization was then conducted on the purified H2A.X-A135C-pS139 at 37 °C for 32 h in the presence of 20 mM TCEP pH 7.5 and glutathione using 10 mM of the radical initiator, 2,2′-azobis[2-(2-imidazolin-2-yl)propane]dihydrochloride (VA-044) (Wako chemicals). The final product i.e., γH2AX containing octamer, was purified by size exclusion chromatography. Samples from each step were analysed by 16% SDS-PAGE. Western blot was performed using rabbit monoclonal anti-H2AX antibody (ab124781) and anti-gamma H2AX (7hosphor S139) antibody (ab81299) to confirm the γH2AX formation.

### TEV digestion

TEV digestion was carried out by adding purified TEV in 25:1 mass ratio and incubating the samples at 4 °C for overnight. The digestion was confirmed by SDS-PAGE. TEV-digested histones were then concentrated up to 4 mg/mL with ultrafiltration using Amicon YM50 membrane (MWCO 50 kDa) at 4 °C (Millipore). The concentrated sample was then injected onto a Superdex 200 10/300GL column. The histone octamer peak was eluted at an elution volume of 12.5 mL. The peak fractions were pooled and concentrated up to 2 mg/ml, aliquoted and flash-frozen in the presence of 50% glycerol for long-term storage.

### 601 – 145 bp DNA preparation

The large-scale DNA purification was performed as described[18]. Briefly, the 145 bp Widom-601 DNA fragment was released from the pUC57 plasmid containing 6 inserts of the 601-145 bp sequence, by EcoRV followed by PEG precipitation. The purity of the 601 DNA as analyzed at each step using 1.5 % agarose gel. After a final size exclusion chromatography step the 601-145 bp DNA was concentrated to 1 mg/ml for nucleosome assembly.

### Nucleosome reconstitution

Nucleosome reconstitution was done essentially as described[35]. The histone octamer and DNA were mixed in the molar ratio of octamer:DNA at 1.1:1.0 and dialyzed sequentially against buffers (10 mM Tris-HCl pH 7.0 and 0.5 TCEP) containing 1 M, 800 mM, 500 mM and 150 mM NaCl, each for at least one hour. The last dialysis was usually done overnight.

### hMDC1 BRCT purification

Hexahistidine tagged hMDC1 BRCT (residues: 1891 – 2089) was cloned into pET47b vector and expressed in Rosetta™ 2(DE3) cell lines (Novagen). Cells in 1L Luria Broth (LB) with 50 mg/ml Kanamycin and 25 mg/ml Chloramphenicol) grew at 37 °C till OD260 reached 0.8. Over-expression was induced by the addition 0.5 mM IPTG and cells continued to grow at 18 °C overnight. Cells were harvested and resuspended in buffer containing 20 mM Hepes pH 7.5, 500 mM NaCl, 1 mM TCEP and protease inhibitor (Roche). The resuspended cells were lysed using Emulsiflex and clarified using centrifugation at 12,000 rpm for 1 hr. The supernatant was incubated with 1 ml of Ni-NTA agarose beads (Qiagen), equilibrated in the lysis buffer for 1 hr. The solution was poured into a gravity column. Beads were then washed with lysis buffer and protein was eluted using a stepwise increase in imidazole concentration (100 mM, 250 mM, 500 mM) added to the lysis buffer. Size exclusion chromatography (SEC) was performed on the Ni-NTA purified protein in a SEC buffer containing 25 mM Hepes pH 7.5, 150 mM NaCl and 1 mM TCEP.

### Fluorescence polarizarion

The purified his – tagged hMDC1 BRCT was tested for its binding affinity with fluorescein labelled γH2AX mimicking peptide (^NH2^-KKCTQApSQEY-^COOH^). A concentration of 50 nM of labeled peptide was used in a reaction volume of 20 μl. Fluorescein fluorescence was excited at a wavelength of 485 nm and the emission was measured at 538 nm on a PerkinElmer Envision plate reader. The change in polarization was graphed as a function of the log of the protein concentration, and the dissociation constant (K_D_) was obtained from the resulting sigmoidal curve.

### Electrophoretic mobility shift assay

The γH2AX containing nucleosome reconstitution and hMDC1 BRCT interaction with these nucleosomes were verified on 5% native polyacrylamide gels run in 1x TBE (Tris-borate-EDTA) at 130 V for 2 h in the cold room. To validate BRCT – nucleosome interaction, 0.1 μM of nucleosome was incubated with increasing concentrations of BRCT in EMSA buffer (20 mM Hepes pH 7.5, 150 mM NaCl, 1 mM MgCl_2_, and 1 mM TCEP). The gel was stained with SYBR Safe DNA gel stain (Thermo Fisher Scientific) to visualize DNA bands. Unmodified nucleosome was used as control.

### Microscale thermophoresis

For the MST experiments, a concentration series of γH2AX containing nucleosome was prepared using a 1:1 serial dilution in buffer supplemented with 20 mM Hepes (pH 7.5), 150 mM NaCl, 1 mM MgCl_2_, 1 mM DTT, and 0.05% Tween 20. The range of nucleosome concentration was from 2.5 μM to a final 0.1 nM, over 12 serial diluted Monolith NT.115 premium capillaries (NanoTemper) with 10 μl samples. The BRCT was labeled using his-tag labeling kit RED-tris-NTA 2^nd^ generation. The interaction for MST experiments was initiated by the addition of 10 μl of 20 nM His tag – labeled BRCT. The measurements were performed on a Monolith NT.115 (NanoTemper) using standard capillaries at 25 °C with 80% MST power, high LED power. Data were analyzed by MO.Control and MO.Affinity software (NanoTemper). The experiments were performed in triplicate.

### Cryo-EM grid preparation and data processing

Three microlitres of γH2AX containing nucleosome samples with or without hMDC1 BRCT at a concentration of 2 μM and were applied on 120 s glow discharged (Solarus) Quantifoil R2/2 gold grids (300 mesh) and flash frozen in liquid ethane on an FEI Vitrobot, using a 0 – 5 s blot time and blotting force of 5 at 4 °C and 100% humidity. CryoEM data were collected using a Titan Krios G2 (Thermo Fisher) operated at 300keV, with a Selectris energy filter (Thermo Fisher) equipped with a Falcon 4 direct detector (Thermo Fisher). Movies were acquired in energy filtered mode with an energy slit of 10kV, magnification of 165kx (pixel size of 0.77 angstroms/pixel). The electron beam had a flux rate of 4.9 electrons/(angstrom)^2^/s, and movies were acquired at 4 frames/s for duration of 10 seconds, for a total electron interaction of 50 electrons/(angstrom)^2^. A 50 mm condenser aperture and a 100 mm objective aperture were used while imaging. The zero-tilt collection was collected with EPU (Thermo) and the tilted collection was performed using Serial EM automated acquisition software. Cryosparc version 4.2.1 was used for the cryo-EM data processing[36]. The 15000 movies obtained were motion-corrected using cryoSPARC’s Patch motion correction (multi). Defocus values were calculated using Patch CTF estimation (multi), and the classes obtained were visually inspected and curated to remove ice contamination, aggregation, and other false positives. 2,512,360 particles were blob-picked using a sphere with a radius of 50 to 200 Å and extracted in a 256-pixel box. 3D volumes of the 2D classified particles were generated using ab initio reconstruction. Several iterations of 2D and 3D classifications were done to remove junk particles and obtain clean 3D reconstructions (Extended Data Fig. 4). For the 601-145 bp DNA and the core histones, the cryo-EM structure of the Xenopus canonical nucleosome (PDB 7OHC)[23] was used. The individual residues were altered to correspond to the sequences of human H2A, H2B, and H4 and performed a rigid body fit using ChimeraX[37]. We built the nucleosome model into the map using iterative rounds of model building and local refinement in Coot[38] and global real space refinement in Phenix[39] leading to the final models (Supplemental Table S1). Our canonical γH2AX nucleosome structure was used to build the stack nucleosome model. The N-terminus of H4 linker conformation adjustment was aided by ISOLDE[40]. Graphics were generated in ChimeraX and PyMol[41]. Local resolution was plotted with Resmap[42].

## Data availability

The cryo-EM maps have been deposited in the Electron Microscopy Data Bank (EMDB) under accession codes EMD-40648 (γH2AX containing canonical nucleosome), EMD-40649 (γH2AX containing nucleosome stack). The model coordinates for nucleosomes were deposited in the Protein Data Bank (PDB) under the accession code 8S06 and 8S07 for the canonical and stack respectively.

## Acknowledgment

This research project was supported in part by High Resolution Macromolecular Cryo-Electron Microscopy facility, University of British Columbia, Canada.

## Author contributions

RP and JNMG conceptualized and designed the study. RP prepared samples and collected the data for Cryo-EM studies. RP and RE processed the Cryo-EM data, carried out model building and Refinement. RP and MTI performed the cloning and construct preparation of the unmodified histone constructs. JL prepared γH2AX construct. RP purified 601-145bp dsDNA. RP and AK performed the western blots. RP and THK performed the FP studies. RP performed the MST and EMSA studies. RP and JNMG wrote the paper with the help and input of all authors.

