## Supplementary data for "Structural Insights into γH2Ax containing Nucleosomes"

**SUPPLEMENTARY FIGURES**

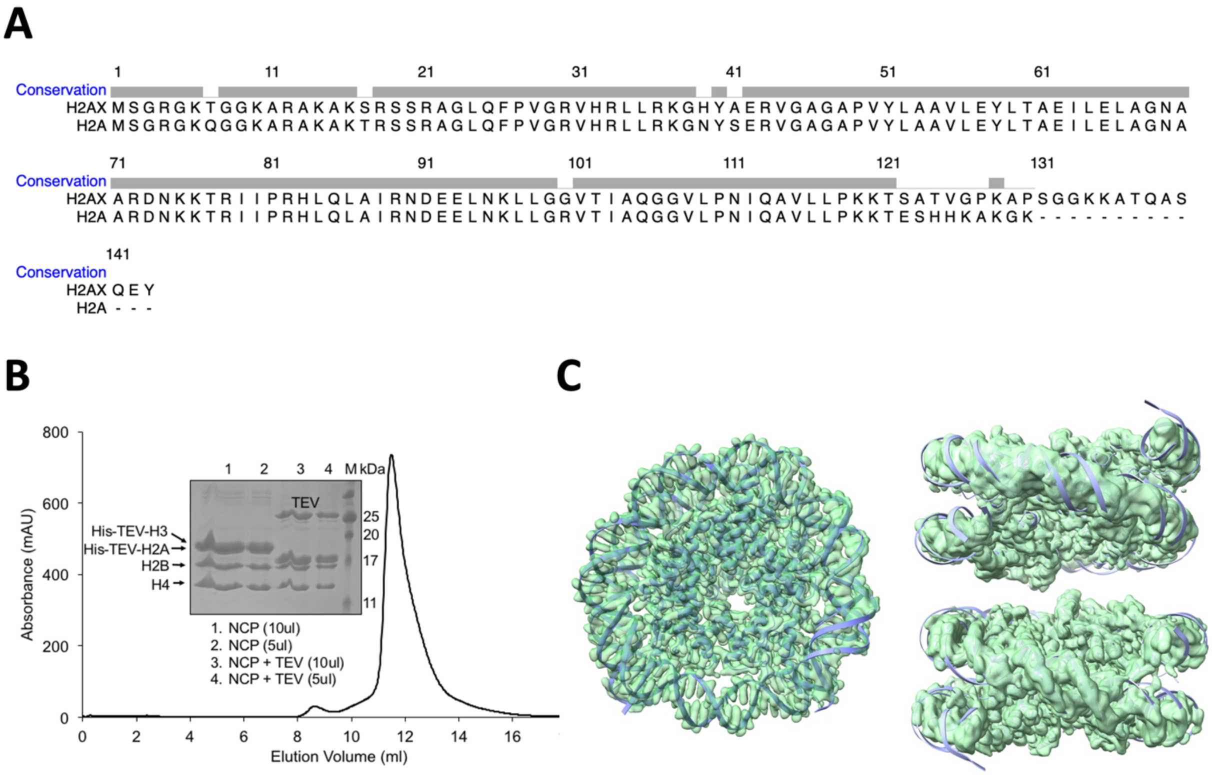

**Extended Data Fig. 1.** Preparation of unmodified H2AX containing nucleosome core particle (NCP) octamer A. Sequence conservation between H2A and H2AX; B. Size exclusion chromatography profile for NCP after the histag had been cleaved with TEV protease. C. Cryo-EM map and model for unmodified H2AX nucleosome (left) and parallel stack nucleosome (right).

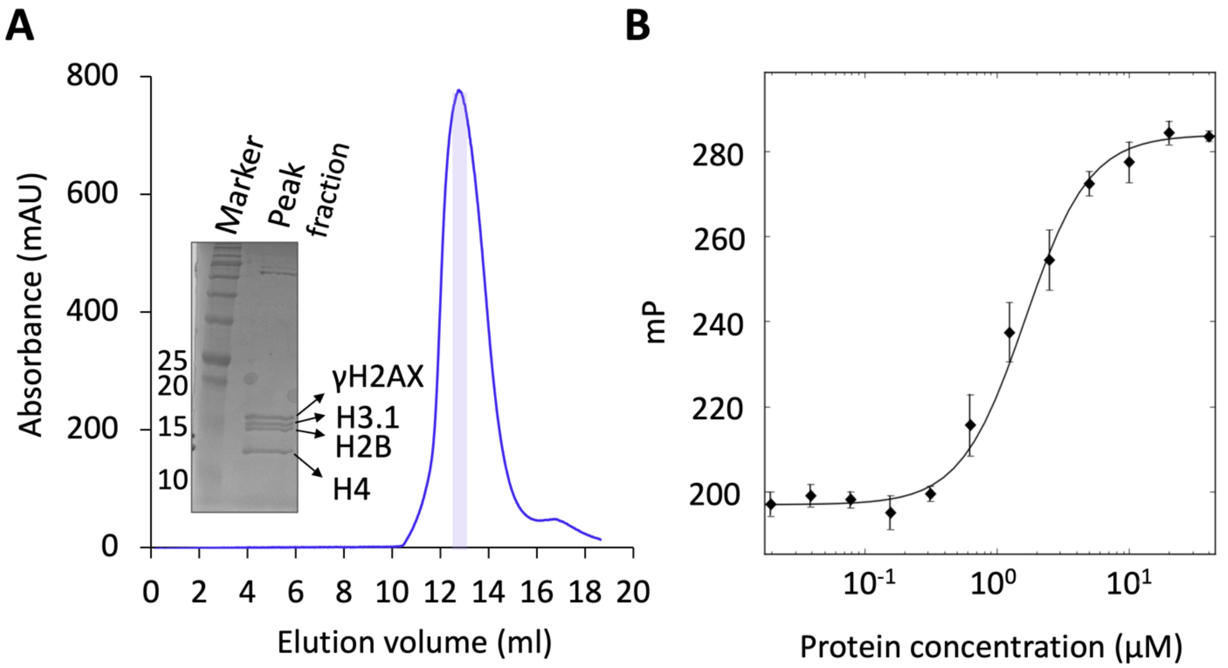

**Extended Data Fig. 2.** Analysis of γH2AX. A. Size exclusion chromatography of γH2AX octamer. The peak fraction in purple was loaded on a 15% SDS PAGE, shown as inset. B. FP studies to validate binding of Fluorescein labeled C-terminal γH2AX peptide with hMDC1 BRCT. The binding affinity is 1.64 +/- 0.14 µM.

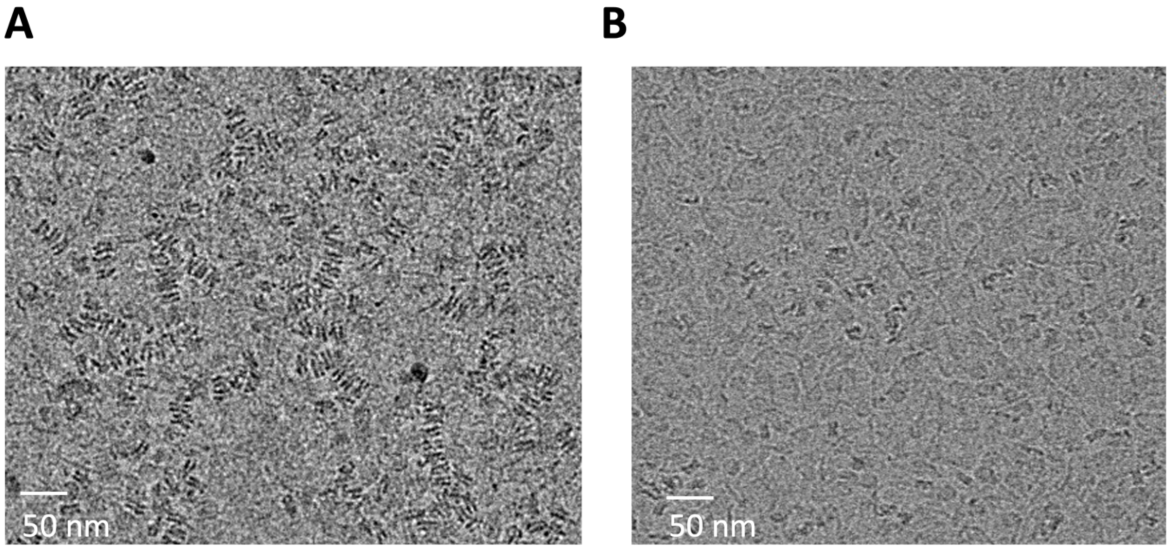

**Extended Data Fig. 3.** Cryo-EM grid images for A. γH2AX nucleosome samples and B. γH2AX nucleosome – BRCT samples. Presence of BRCT inhibits the inter-nucleosomal stacking interactions.

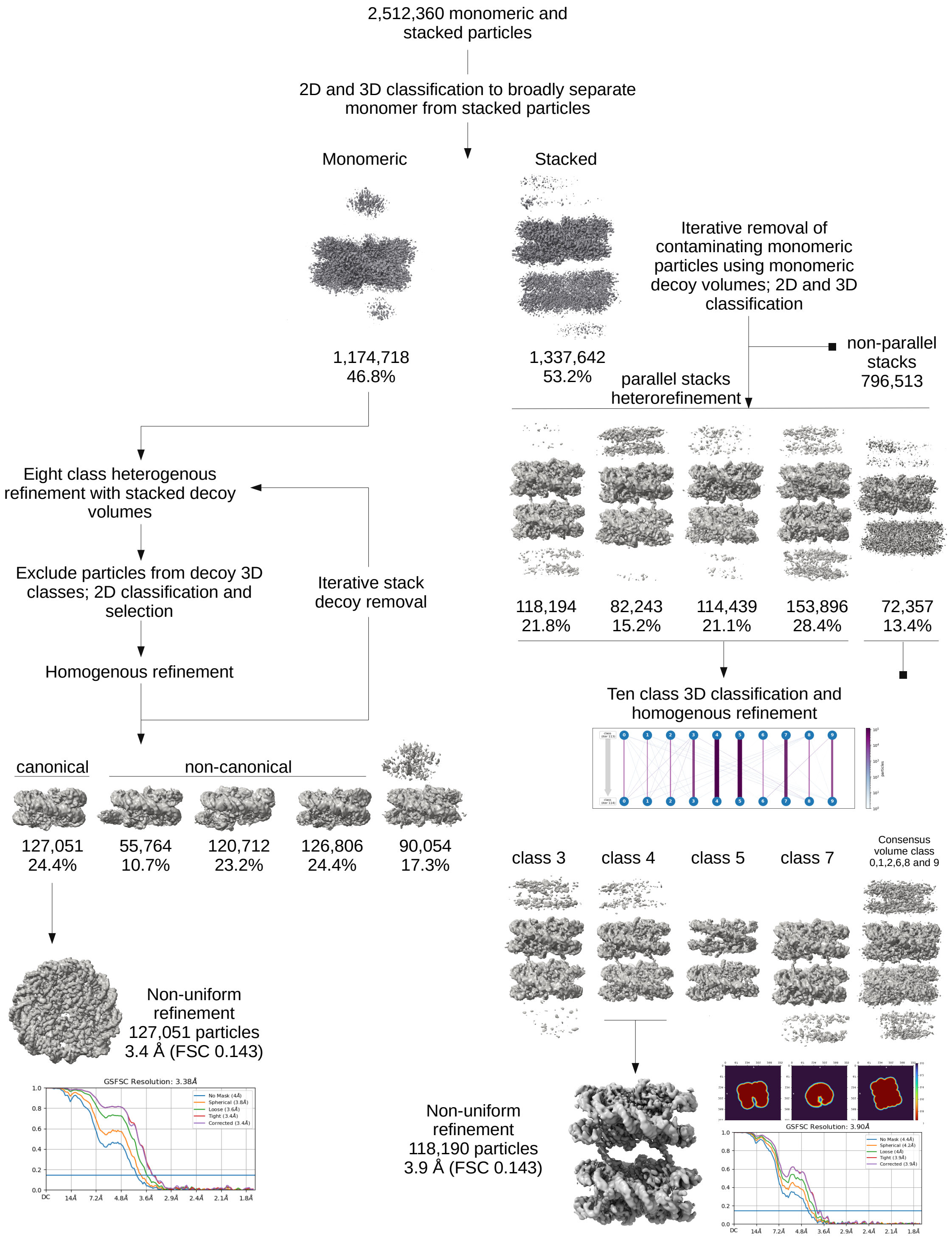

**Extended Data Fig. 4.** Flowchart for cryo-EM data processing for γH2AX containing nucleosome as described in the Methods.

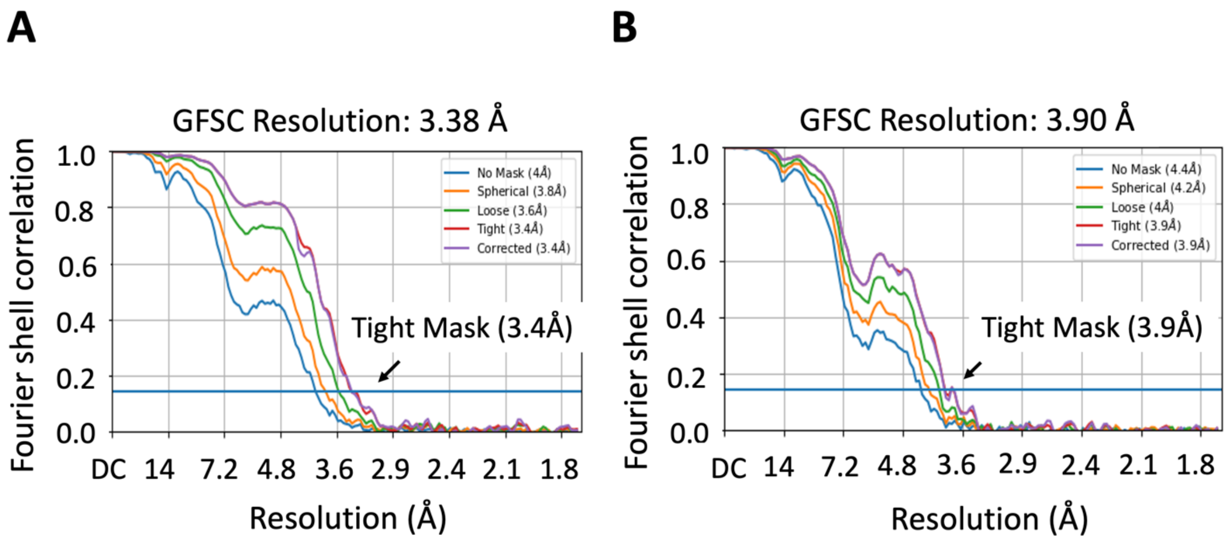

**Extended Data Fig. 5.** Gold standard Fourier shell correlation (GFSC) curve. A. Canonical γH2AX containing nucleosome B. Stack γH2AX containing nucleosome. The resolution reported here was according to the 0.143 criterion indicated by the blue line.

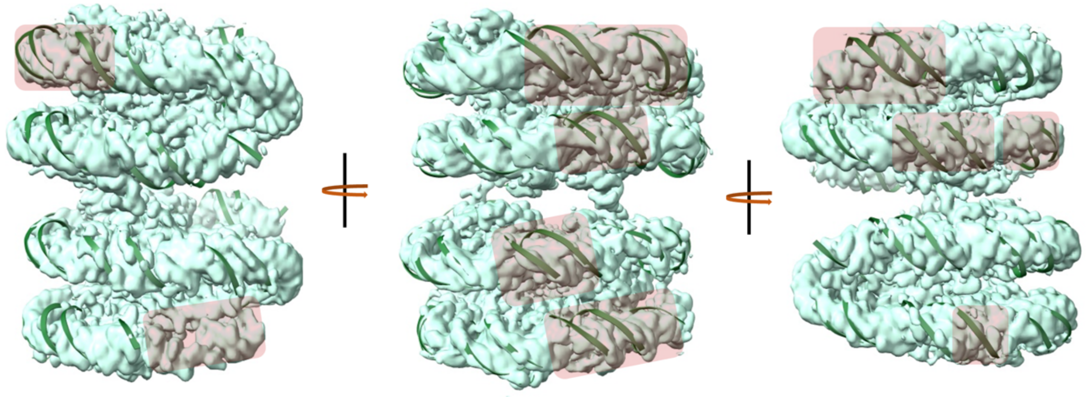

**Extended Data Fig. 6.** DNA backbone flexibility observed in γH2AX nucleosome stack structure. The nucleosome density is shown in the surface representation and the model is shown in green cartoon. Different views of the structure are shown where the flexible DNA backbone are shaded.

### Supplemental Table S1. Data collection and refinement statistics

| **Data Collection** | | | | |
|---|---|---|---|---|
| Microscope | Titan Krios | | | |
| Magnification | 165000x | | | |
| Voltage (kV) | 300 | | | |
| Angstrom per pixel (Å) | 0.86 | | | |
| Stage tilt (°) | | 0 | 17 | 34 |
| Imaging software | | TEM | SerialEM | SerialEM |
| Dose per frame (e/Å$^2$) | | 1.25 | 1.25 | 1.26 |
| Frame duration (s/frame) | | 0.05 | 0.06 | 0.04 |
| Exposure time (s) | | 2 | 2.3 | 1.6 |
| Electron exposure (e/Å$^2$) | | 50 | 50 | 50.23 |
| Number of movies | | 7289 | 5102 | 11283 |
| Defocus range (µm) | 1.5-2.1 | | | |
| Nucleosome Particle Type | *Canonical* | *Parallel stack* | | |
| Symmetry imposed | C1 | C2 | | |
| Final particles | 127051 | 118190 | | |
| Map resolution (FSC 0.143, Å) | 3.4 | 4.0 | | |
| CC (volume) | 0.74 | 0.5 | | |

| **Refinement and Validation** | | |
|---|---|---|
| Nucleosome Particle Type | *Canonical* | *Parallel stack* |
| Composition (#) | | |
| Chains | 10 | 20 |
| Atoms (hydrogens) | 21257 (9457) | 40981 (18342) |
| Residues, protein | 751 | 1488 |
| Residues, nucleotide | 286 | 530 |
| Bonds (RMSD) | | |
| Length (Å) (# > 4σ) | 0.006 (0) | 0.005 (0) |
| Angles (°) (# > 4σ) | 0.989 (2) | 0.938 (5) |
| MolProbity score | 1.22 | 1.16 |
| Clash score | 4.43 | 3.65 |
| Ramachandran plot (%) | | |
| Outliers | 0.00 | 0.07 |
| Allowed | 1.36 | 1.85 |
| Favored | 98.64 | 98.08 |
| Rotamer Outliers (%) | 0.00 | 0.81 |
| Cβ outliers (%) | 0.00 | 0.00 |
